# Structure of Topologically Associating Domains and gene regulation embedded in the 5C data revealed by a polymer model and stochastic simulations

**DOI:** 10.1101/065102

**Authors:** O. Shukron, D. Holcman

## Abstract

Chromatin organization is probed by chromosomal capture data, from which the encounter probability (EP) between genomic sites is represented in a large matrix. However, this matrix is obtained by averaging the EP over cell population, where diagonal blocks called TADs, contains hidden information about sub-chromatin organization. Our aim here is to elucidate the relationship between TADs structure and gene regulation. For this end, we reconstruct the chromatin dynamics from the EP matrix using polymer model and explore the transient properties, constrained by the statistics of the data. To construct the polymer, we use the EP decay in two steps: first, to account for TADs, we introduce random connectors inside a restricted region defining the TADs. Second, we account for long-range frequent specific genomic interactions in the polymer architecture. Finally, stochastic simulations show that only a small number of randomly placed connectors are required to reproduce the EP of TADs, and allow us to compute the mean first time and the conditional encounter probability of three key genomic sites to meet. These encounter times reveal how chromatin can self-regulate. The present polymer construction is generic and can be used to study steady-state and transient properties of chromatin constrained on 5C data.

## 1 Introduction

The chromatin molecule is organized in heterogenous sub-regions of various sizes, as recently revealed by Chromosome Capture (5C) data [1, 2]. This multi-scale organization is due to short and long-range genomic interactions between DNA segments, collected over large cell population. It was recently shown [3, 4] that the mammalian chromatin, at a resolution of 3kB, contains an organization at 1Mbps scale, where some structures are enriched in intra-connectivity, reflecting an increased encounter probability between these segments. These encounter properties are represented in a two-dimensional encounter frequency (EF) matrix, containing diagonal blocks called Topologically Associating Domains (TADs) [3, 5]. These blocks associated with gene regulation [3], DNA replication units [6], or DNA entanglement and cross-linking by gluing molecules such as cohesin, CTCF, and condensin [3].

Looping between chromatin sites are precisely the events sampled by chromosome capture data (3C, 4C, 5C, HiC) [3, 7], and single cell HiC confirms that configurations can vary between cell types and phases [8]. In that context, TADs represent averaged chromatin conformations, characterized by a higher mean number binding compared to non-TAD regions.

To interpret the encounter probability (EP) in the 5C data, polymer models are utilized, such as the Rouse polymer [23] that predicted a -3/2 decay exponent for the EP with genomic distance. Polymer models have been used to described chromosomal territories [9], further suggesting that inactive genes are located inside these territories. By adding local binding sites on polymer models, TAD structures can emerge [10, 11, 12, 13], where the decay exponent of the EP can vary at different scales. 5C EPs at the X-inactivation centre was recently used to calibrate interaction potentials between beads in a freely jointed chain, allowing to assess internal 3D mean distances inside TADs [5]. By incorporating reversible binding of diffusing molecules on a self-avoiding polymer [14], some chromatin conformation have been identified. Inter chromosome distances at large scale resolution of 1Mb pairs were also found using minimization algorithms [15]. In parallel, single particle trajectories of tagged DNA locus [16, 17, 18, 19, 20, 21, 22] reveal that chromatin is constantly remodeled, characterized by mean square displacement function and its anomalous behavior.

Our aim here is to elucidate the relationship between TADs structure and gene regulation. For this end, we present a general procedure to construct a coarse-grained polymer model that accounts for the statistical properties of the chromosome capture data, such as the decay of the EP and TADs. The reconstructed polymer is then used to study transient properties such the interaction between two given sites. We start with a Rouse polymer model, which consists of beads connected by harmonic spring, and use the EP of the 5C data of mammalian X chromosomes to constrain further monomer interactions. The construction procedure is divided into two steps: first, to account for heterogeneity in the 5C data, we add connectors (cross-links) between genomic sites chosen at random, and show we can reproduce TAD block. We show that the number of random connectors that is needed to be added is uniquely determined from data. However, this step is insufficient to recover the EP decay probability. Thus, in the second step, we account for consistent long-range interaction, represented by the local maximum of the EP matrix. We validate this reconstruction by showing that the EP-matrix, constructed from simulations of the polymer following the two-step procedure, has the same decay exponent (for each monomer) as that of the empirical data. Once the polymer model is constructed, we estimate the conditional encounter probability and the associated mean first passage time between three specific genomic sites. The present polymer construction is generic and can be used to study transient properties inside or between any TADs or any 5C matrix.

## 2 Materials and Methods

### 2.1 Construction a generalized Rouse polymer from encounter frequency map

The Rouse model describes a polymer as a collection of beads *R*_*n*_(*n* = 1…*N*) connected by harmonic springs and driven by Brownian motion. The energy of the polymer is given by [23]

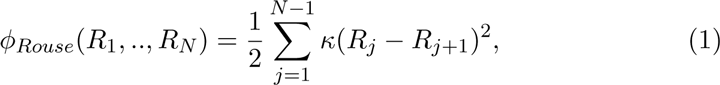

where here we set 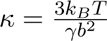 and *b* is the standard deviation of the distance between adjacent monomers, γ is the friction coefficient, *k*_*B*_ the Boltzmann coefficient and *T* the temperature.

To account for a sub-chromatin region, characterized by a higher encounter probability than the rest, we added connections between monomer-pairs chosen randomly (with uniform distribution) inside this subregion 𝒞_𝒩_ such that an additional potential

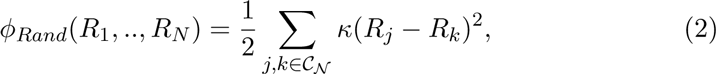

is added to *ϕ_Rouse_*, where 𝒞_𝒩_ is the ensemble of indices defining the regions. In addition, to account for consistent long-range interactions, reflected by peaks in EP matrix (Fig. 1C), we connected monomer-pairs by spring as follows: first, the positions of peaks form a subset *S*_*Max*_ of the ensemble of the off-diagonal local maxima in the EP-matrix, such that their EP is higher than a threshold *T*_*th*_. In practice, we chose the threshold *T*_*th*_ to represent the encounter probability for the nearest neighbor monomers in *M*_*i,j*_, that is

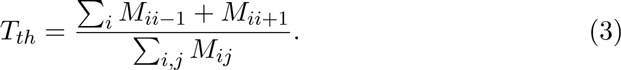

**Figure 1:**
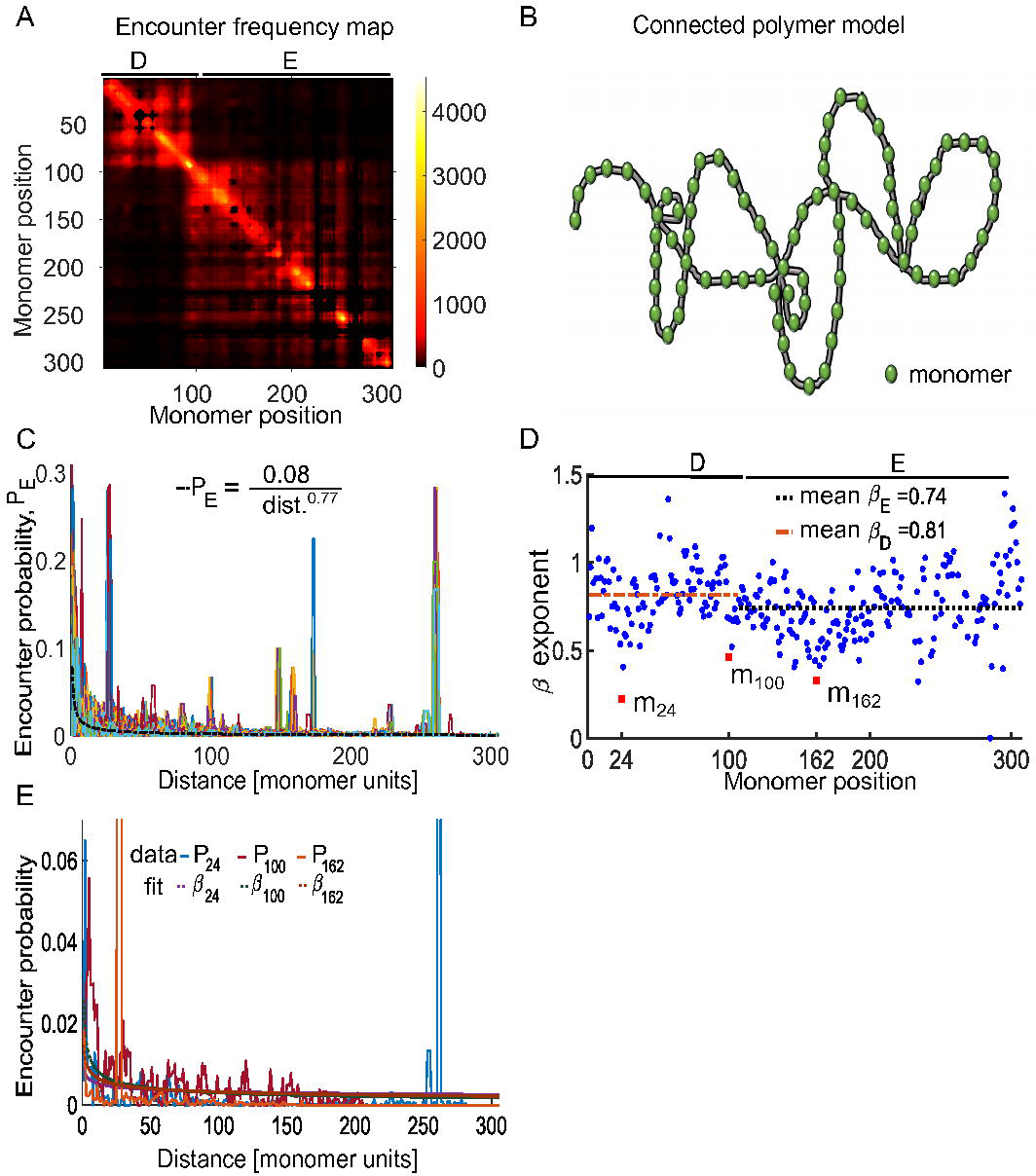
Statistics of Conformation Capture data. **A.** Average encounter frequency map of two 5C replica spanning ≈ 1Mbp genomic region containing two Topologically Associating Domain (TAD) D (monomers 1-106) and TAD E (monomers 107-307) [3], where the map was coarse-grained into 307 monomers of size 3kb [5]. **B.** Schematic representation of a polymer model with randomly connected monomers. **C.** Empirical encounter probability, *P*_*n*_, for monomer *n* plotted with respect to the genomic distance *d* [monomer units], reveals long-range interactions (localized peaks). *P*_*n*_ are fitted with functions *Ad*^−β^, where *β* is the decay exponent and *A* the normalization factor. For the mean encounter probability 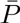, we have *A* = 0.08 and *β* = 0.77 (thick red curve). **D.** Distribution of the β_*n*_ exponents (*n* = 1‥307) (blue dots) displaying high variability: Monomers *m*_24_, *m*_100_, *m*_162_ (red square dots) with β_24_ = 0.22, β_100_ = 0.46, and β_162_ = 0.33, respectively, accounts for high peaks (first and last), while the middle one corresponds to the boundary between TADs. **E.** Encounter probability *P*_*n*_ for monomers n=24, 100, and 162, corresponding to local minima shown in box D.

The spring constants *κ*_*m,n*_ between monomer *m* and *n* in *S*_*Max*_ are determined from the empirical encounter probability *P*_*m,n*_ at each peak position. For a Rouse chain the joint probability density function of beads *R*_*m*_ and *R*_*n*_ is given by [23, p.15]

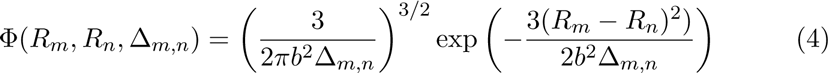

where Δ_*m,n*_ = |*m* − *n*|. For the nearest neighbors Δ_*m,n*_ = 1 and the encounter probability occurs at small distances so that the exponential is almost 1.

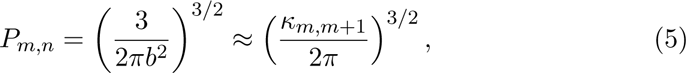

We approximate the chromatin as a polymer chain with a uniform variance *b* ^2^ between adjacent monomers, thus, the constant 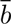 is estimated as the mean EP over all neighboring monomers:

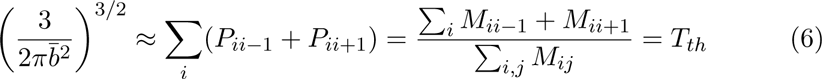

To account for the long-range interactions, we applied formula 5 to estimate the effective spring constant from the empirical EP 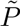,

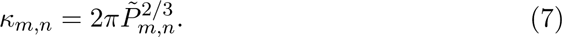

The energy related to peak interactions is described by

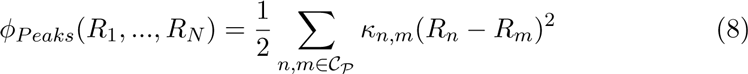

where 𝒞_𝒫_ is the ensemble of monomer indices associated to the selected peak positions. In summary, the total energy of a polymer containing random connected and prescribed peaks, is the sum of three energies 1-2,8,

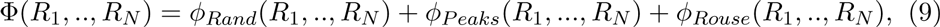

and the stochastic equation of motion for *n* = 1, ‥,*N* is

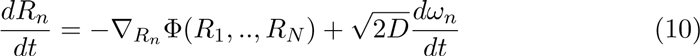

where 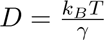 is the diffusion constant, γ is the friction coefficient, and *ω*_*n*_ are independent 3-dimensional Gaussian noise with mean 0 and standard deviation 1.

### 2.2 Polymer model associated 5C data

To account for the 5C-data, comprised of a subsection of the X-chromosome from female mice embryonic stem cells reported in [3], showing TAD D and E as two diagonal blocks (Fig. 1A), we use the coarse-grained procedure of [5], with a polymer of length *N* = 307. Each monomer represents a genomic segment of 3*kb* and is connected to its 2 nearest neighbors by an harmonic spring (see subsection above). TAD D (resp. E) is represented by the range of beads from 1 to *N*_*D*_ = 106 (resp. 107 to 307). The number of monomer is TAD E is *N*_*E*_ = 201.

To reproduce the empirical EP extracted from data, we connected non nearest neighbor pairs of monomers chosen randomly with probability 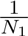 (resp. 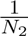)) where *N*_1_ = (*N*_*D*_ − 2)(*N*_*D*_ − 1)/2 (resp. *N*_2_ = (*N*_*E*_ − 2)(*N*_*E*_ − 1)/2). The number of connectors in each TAD is a fraction *ξ* of the total possible number of non nearest neighbor pairs, for TAD D (resp. E), we have 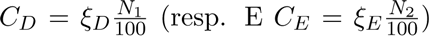 and ξ_*D*_, ξ_*E*_ ∈ [0, 100] will be extracted from data. Random connectors were not added between monomers belonging to different TADs. The procedure of adding random loops in a Rouse polymer is implemented using the energy of random loops, as described in the previous section. Finally, 24 connectors were added (see SI) to all polymers, corresponding to the selected peaks present in the EP matrix.

### 2.3 Numerical simulations of the reconstructed polymer model

Using the method described in the two previous subsections, we generated polymer realizations, each differ in the position of random connectors inside each TAD. To generated statistics for the EP, we simulate the polymer past its relaxation time τ_*R*_. The time τ_*R*_ is determined for each realization by τ_*R*_ = 1/(*κ*_min_ λ_1_), where *D* is the diffusion coefficient, *κ*_*min*_ is the minimal positive spring constant, and λ_1_ is the smallest non-vanishing eigenvalue of the polymer’s connectivity matrix [24], which we calculate numerically. In simulations, we divide equation 10 by 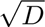 and the spring constants are scaled by the fiction coefficient such that 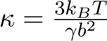. The encounter frequency matrix of the 307 monomers is computed at time τ_*R*_, where two monomers are considered to have encountered if their distance is less than ∈. The time step for all simulations is Δ*t* = 10^−2^[*sec*].

#### 2.3.1 Parameters

We summarize in Table 1 the values of parameters used in simulations

**Table 1:**
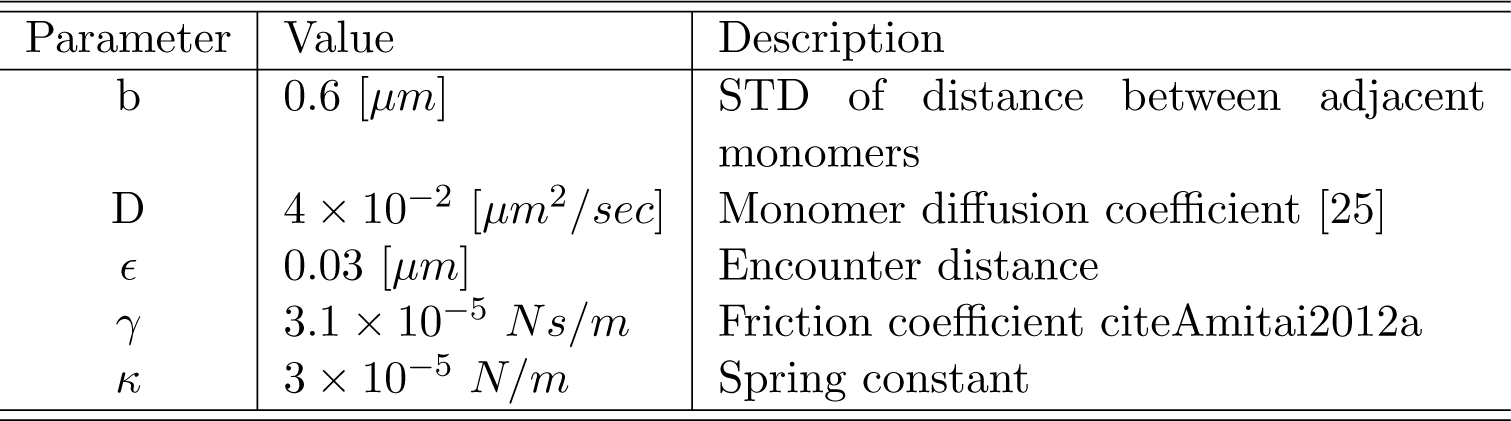
values of simulation parameters

| Parameter | Value | Description |
| --- | --- | --- |
| b | 0.6 [$\mu m$] | STD of distance between adjacent monomers |
| D | $4 \times 10^{-2}$ [$\mu m^2/sec$] | Monomer diffusion coefficient [25] |
| $\epsilon$ | 0.03 [$\mu m$] | Encounter distance |
| $\gamma$ | $3.1 \times 10^{-5}$ $Ns/m$ | Friction coefficient citeAmitai2012a |
| $\kappa$ | $3 \times 10^{-5}$ $N/m$ | Spring constant |

## 3 Results

### 3.1 The encounter probability of coarse-grained 5C data

We first described how we constructed a polymer model from the symmetrized 5C frequency matrix *M* (Fig. 1A). By symmetrizing the EP matrix, we averaged-out asymmetrical fluctuations. The 5C data we used represent a sub-region of the X chromosome (≈ 92,000 bps), that was previously segmented into two regions called Topological Associating Domains (TADs) D and E [3]. The matrix *M* was further coarse-grained by beaning the encounter frequencies into 307 monomers of 3*kbps* [5], where TAD D (resp. TAD E) is represented by the first 106 monomers (resp. 107-307), as shown in Fig. 1A. We introduce a general polymer model (Fig. 1B) with arbitrary configuration, the properties of which will be extracted from *M*_*em*_.

The encounter probabilities between monomer *m* and monomer *n* are computed from matrix *M* by

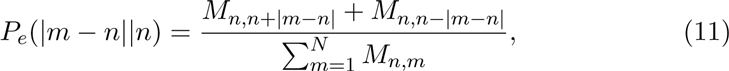

which depends on the genomic distance |*m* − *n*| (Fig.1C). Although the encounter probabilities decayed with |*m*−*n*| for each *n*, they contain peaks that reflect consistent long-range interactions between monomers. To quantify the decay of the EP, we fitted its average value 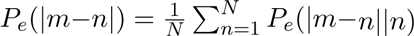 (black dotted line in Fig. 1C) with the function

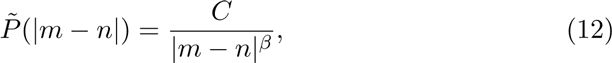

where *C* and *β* > 0 are two constants. For a Rouse polymer, the EP function 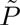 is characterized by a decay exponent *β* = 3/2 [23]. Fitting (12) to data, revealed that *β* = 0.77, from which we concluded that the polymer model should be modified to account for higher compaction than allowed by a Rouse polymer [26].

To better account for the heterogeneity in the EP of each monomer, we plotted the distributions of the exponent β_*n*_ for *n* = 1..307 along the polymer (Fig. 1D blue dots). The exponents β_*n*_ were extracted by fitting the function 12 to the empirical EPs 11. The large variability in β_*n*_, *n* = 1..307 reflects the local heterogeneity of the chromatin architecture at the current scale (a monomer represents 3kbps). The average *β* for TAD D and E was found to be β_*D*_ = 0.81 and β_*E*_ = 0.78, respectively. The local minima of *β* deviated significantly from the mean values (Fig.1D red squares), which can be associated with chromatin features (Fig. 1E): such values can arise due to specific long-range interactions, or boundary between chromatin subdomains. Point *m*_100_ (monomers 102-107 in Fig. 1D red) is indeed located at the boundary between TAD D and E, while *m*_24_ and *m*_162_ are characterized by strong long-range interactions. To conclude, the distribution of *β* values extracted from the EP is heterogeneous, which can disclose chromatin sub-regions and long-range strong interactions. We shall account in the next two sections for these characteristic features and include in our polymer model both random, and persistent long-range connections between monomer.

### 3.2 Encounter probability of random loop polymer model

To determine the level of connectivity that should be added to a generalized Rouse polymer in order to reproduce the EP-decay with a prescribed exponent β, we first studied the case of one TAD-like region in a 307 monomer chain. We added connections between random non nearest neighbor monomer-pairs in the subregion 103-203 (Fig. 2A). The number of connectors, or the connectivity percentage *ξ* (number of connected monomer-pairs, see Materials and Methods), was increased in the range 0 − 2%. By adding connectors, the EP between distant monomers has increased, as represented in the EP-matrix (Fig. 2B). In contrast, outside the region 103-203 the EPs were similar to the case ξ = 0 (linear chain), showing that the connected region did not affect the EP in the non-connected one. At this stage, we have shown that adding random connectors allows recovering some statistical characteristics of a TAD region.

**Figure 2:**
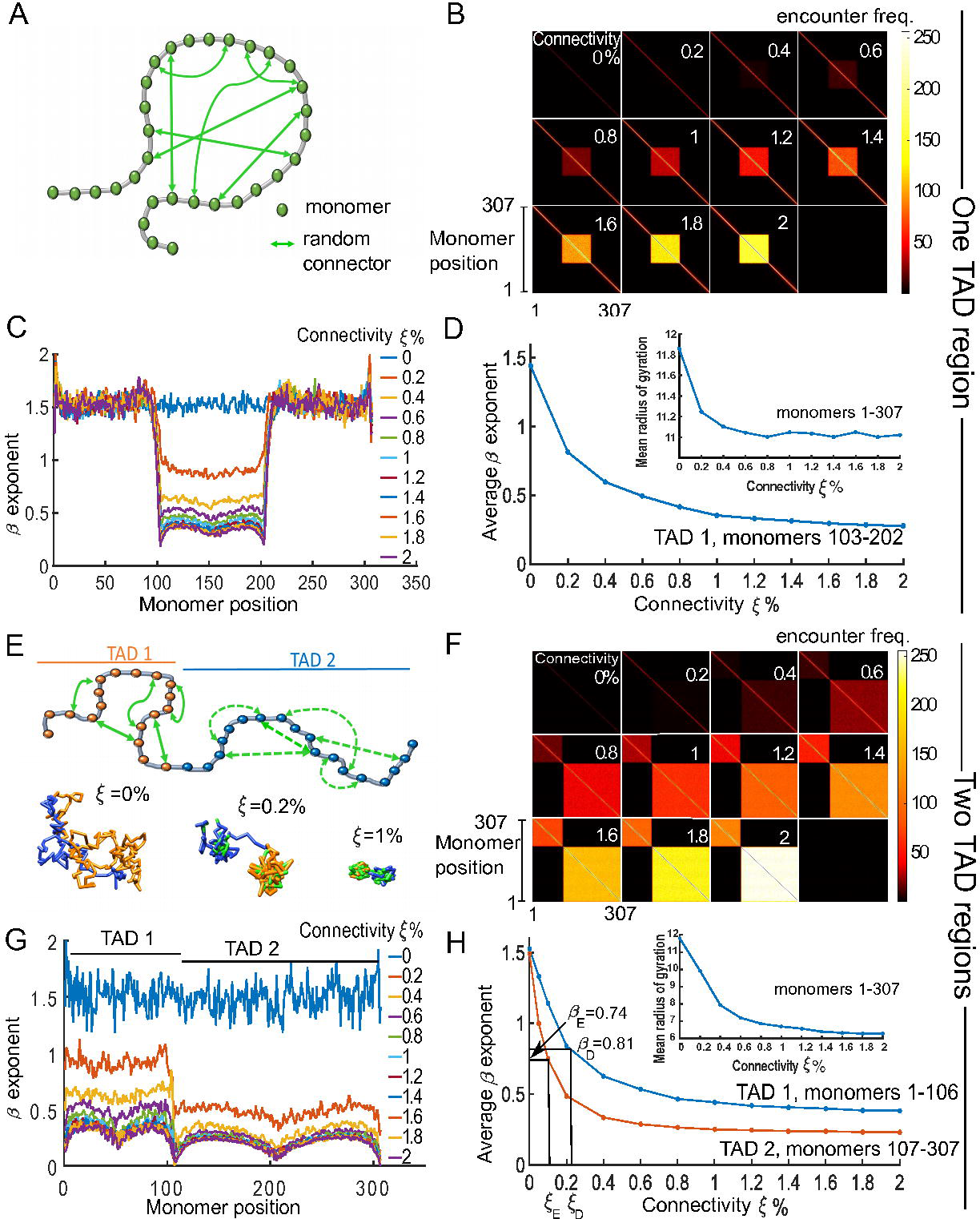
Statistics of simulated generalized Rouse polymer chain for various connectivity in one and two sub-regions. **A.** Schematic bead-spring chain connected at random positions (two-sided green arrows) between non-nearest-neighbor monomers. **B.** Encounter frequency maps of a 307 monomers chain, where connectors are added randomly between monomers 103-202 for each realization. TAD-like structure emerges as the connectivity *ξ* (number of connectors) increase from 0 to 2%. **C.** Distribution of β_*n*_ (*n* = 1‥307) fitted by the function *Ad*^−β^, where *d* is the distance along the chain [monomer units], to the encounter probabilities of numerical simulation. The TAD regions is characterized by a low *β* values. **D.** Average value of *β* for monomers in the interval 103-202 with respect to the connectivity percentage *ξ*, which is decaying due to the increase of long-range interaction between monomers of the polymer. This decay is correlated with the one of the radius of gyration (embedded sub-figure). **E.** Schematic polymer chain, where two defined regions: monomers 1-106 (TAD 1, orange circles) and monomers 107-307 (TAD 2, blue circles), are randomly connected (green arrows). No connections were added between the two TAD regions. We present in the lower panel, three snapshot realizations of a random loop chain with TAD 1 (orange) and TAD 2 (blue) and random connectors (green) for three increasing values of connectivity *ξ* = 0, 0.2, 1%. **F.** Encounter frequency maps showing the two TAD regions for an increasing number of random connectors. **G.** Distribution of *β* exponent for ξ ∈ [0, 2]%, showing the border (n=106) effect between TADs. **H.** Average *β* over each TAD 1 (blue) and TAD 2 (orange) for *xi* ∈ [0, 2]. The curves decrease until plateau at 0.42 (0.24) for TAD 1 (resp. TAD 2). We use these curves to recover the connectivity percentage *ξ* of the experimental TAD D, with β_*D*_ = 0.74 (resp. TAD E with β_*E*_ = 0.81) for which ξ_*D*_ = 0.23% (resp. ξ_*E*_ = 0.12%.

To find the optimal number of connectors necessary to recover a given TAD, we set out to elucidate the relationship between the connectivity percentage *ξ* and the decay exponent β. We have simulated an ensemble of polymer to their relaxation time (see Materials and Methods), and used the equilibrium configuration to estimate the EP of each monomer for *ξ* ∈ [0, 2]%. We calculated *β* by fitting the function 12 to simulation data: the values of β_*n*_ for *n* ∈ [103, 203] decreased with *ξ*. Indeed, for *ξ* ≈ 0.2%, the coefficients β_*n*_ decreased below the Rouse exponent (equal to β_*Rouse*_ = 1.5), indicating compact polymer configurations. For *ξ* = 2%, the mean exponent β_*n*_ and *n* = 103 − 203 was 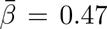, with a minimal value 0.42 obtained for the boundary monomers 103 and 203 (Fig. 2C). These results confirm that the polymer condenses in smaller regions, measured by the mean square radius of gyration *R*_*g*_ [23], decaying from 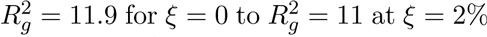 (Fig. 2D). Values of *β* outside the TAD region (monomers 1 to 102 and 204-307) were mostly unchanged, fluctuating around *β* = 1.5, confirming that statistical Rouse properties are not affected when connectors are added to the middle region (103-203). Finally, the average value of β_*n*_ (computed over *n* = 103 − 203) versus *ξ* is shown in Fig. 2D and, as we shall see, can serve to extract the connectivity percentage *ξ* from the empirical data.

To reproduce the two TAD E and D regions of the X-chromosome, we started with a polymer of 307 monomers and added random connectors (green arrows) between monomers 1-106 and between monomers 107-307, as described in Fig. 2E upper panel). This partition follows the empirical TAD segmentation [3, 5]. Three polymer realizations for *ξ* = 0, 0.2, 1% are shown in Fig. 2E bottom panel, showing polymer condensation into two regions. The encounter frequency matrix show that two TAD-like regions, named TAD1 and TAD2 (Fig. 2F), emerge as *ξ* increases from 0 to 2%. To extract the exponent *β* (Fig. 2G, colored curves) we fitted the function 12 to the EP-matrix for each monomer inside TAD1 and 2. In both cases, the exponent *β* decreased below β_*Rouse*_ = 1.5 (*ξ* = 0% blue curve) and for *ξ* = 0.2%, 9 and 39 random connectors were added for TAD1 and TAD2, respectively. The boundary between TADs is characterized by an abrupt decay of the beta value. The *β* exponent (averaged over each TAD), plotted with respect to the connectivity *ξ* (Fig. 2H), was used to determine the number of connectors necessary to reconstruct the empirical data. Indeed, we extracted from the EP matrix (Fig. 1A) that *β*_*D*_ = 0.81, β_*E*_ = 0.78 the associated connectivity percentages ξ_*D*_ = 0.12,ξ_*E*_ = 0.23, respectively (Fig. 2H). To conclude, we estimated the number of random connectors needed to be added on a Rouse polymer such that the *β* exponent of the reconstructed and empirical EP-matrix are identical.

### 3.3 Incorporating long-range empirical interactions in the polymer model

Another key feature present in the 5C EF-matrix (Fig. 1A) is the ensemble of consistent long-range interactions between monomers (Fig. 1C). To account for these interactions, we connected monomers corresponding to off-diagonal local maxima of the EF matrix, which exceed a given threshold (see Materials and Methods). We found 24 long-range connections: 7 (resp. 13) inside TAD D (resp. E) and 4 across the two (see SI for the list of monomers pairs) as shown in Fig. 3A. As revealed by simulations of a Rouse polymer with added connectors, the polymer configuration are condensed, characterized by the radius of gyration about R_*g*_ = 9.1 (compared to 12 for the Rouse chain). Three realizations of the polymer are shown in Fig. 3A. We adjusted the spring constant *κ*_*m,n*_ between monomer *m* and *n* corresponding to consistent long range interactions, based on the values of their EPs (see Materials and Methods). The scaled coefficient *κ*_*m,n*_ are summarized in the SI, and are between 1.1 to 3 times higher than the spring constant assigned to connectors of the linear backbone (*κ* = 0.97).

**Figure 3:**
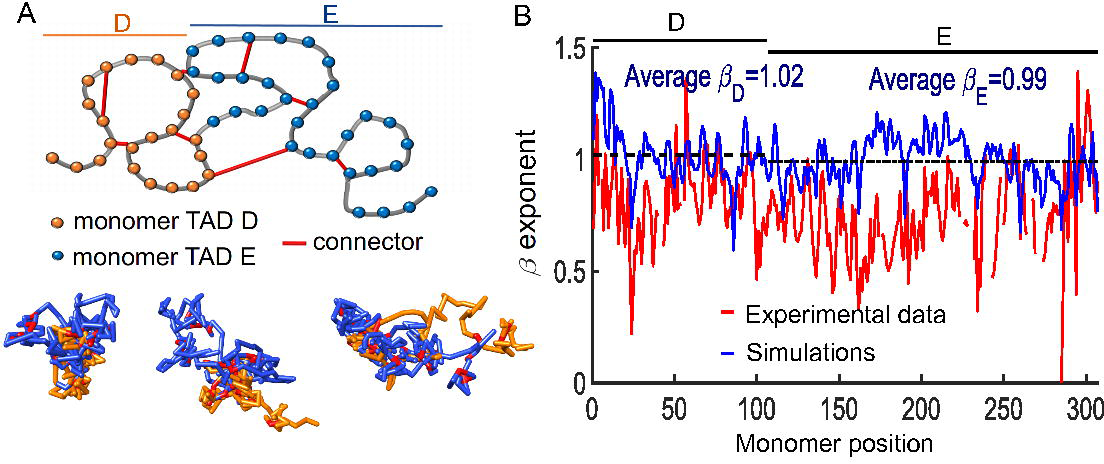
Effect of persistent long-range connectors on polymer folding. The connectors account for peaks present in the empirical 5C encounter frequency maps. **A.** Upper panel: schematic representation of the bead-spring polymer model with added fixed connectors (red) representing specific long-range monomer interactions (peaks) shown in Fig. 1C. Lower panel: three different realization of the same polymer, showing TAD D (orange), TAD E (blue), and fixed connectors (red). **C.** Simulated (blue) and experimental (red) *β* exponent of the fitted encounter probability. The polymer model contains only specific long-range interactions (no random connectors ξ_*D*_ = 0, ξ_*E*_ = 0). Such a model cannot reproduce the encounter frequency maps of the experimental data. The average *β* values for TAD D and E are β_*D*_ = 1.02 (resp. β_*E*_ = 0.99).

To quantify the effect of adding long-range connections, we computed the exponents *β*_*n*_ by fitting the function 12 to the EPs from simulations, and compared it to the ones computed from the experimental data (Fig. 3B). The experimental β_*n*_ (red) were generally lower than the ones obtained from simulations (blue), indicating that the reconstructed chromatin polymer is more condensation in both TADs. Furthermore, we concluded that the addition of long-range connectors is insufficient to reproduce the statistics of the 5C data.

### 3.4 Combination of random loops and long-range interactions to construct a polymer model of a TAD

We previously evaluated separately the effect of random connectors and long-range interactions on the EPs. We computed the decay exponent β, that we compared with coarse-grained 5C data. We now combine these two constraints, such that specific and non-specific connectors are added to a generalized Rouse polymer (Fig. 4). The connectivity percentage matching that of the experimental data in each TAD was found to be ξ_*D*_ = 0.23% and ξ_*E*_ = 0.12%. These value are summarizing the sum of the contribution from the two types of connectors (see Fig.2H). For long-range specific interactions (Fig. 3), we have previously obtained β_*D*_ = 1.02,β_*E*_ = 0.99, corresponding to ξ_*D*_ = 0.12%,ξ_*E*_ = 0.07%(Fig. 2H) for TAD D and E, respectively. We, therefore, attributed the remaining percentages to the addition of random connectors, that is ξ_*D*_ = 0.11,ξ_*E*_ = 0.05. The number of added random connectors corresponding to ξ_*D*_ = 0.11%,ξ_*E*_ = 0.05% are 6 and 10 in TAD D and E, respectively.

**Figure 4:**
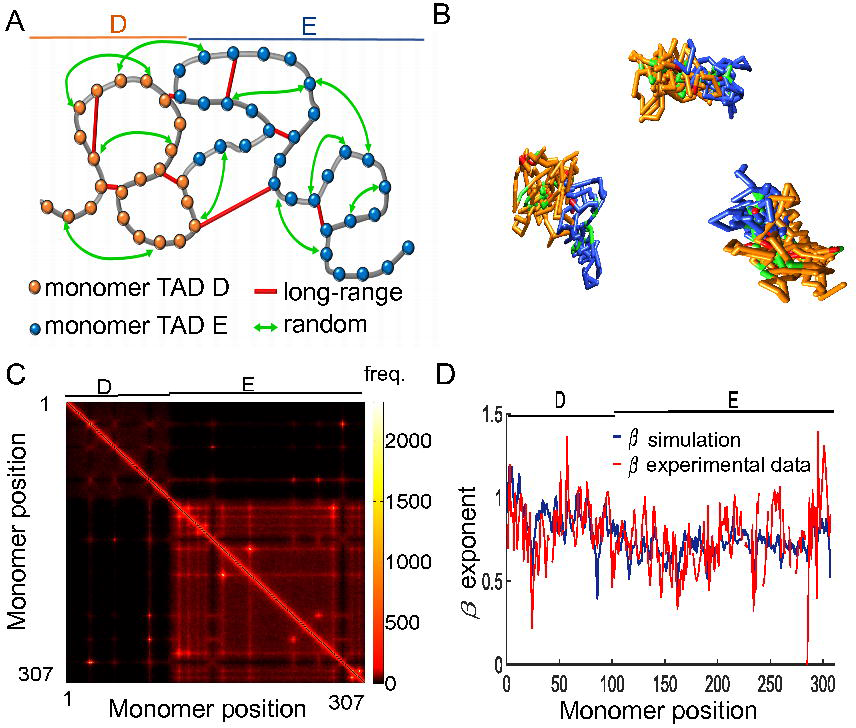
Coarse-grained reconstruction of chromatin using extracted random loops and connectors corresponding to peaks of the 5C data. **A.** Schematic polymer model, where TAD D (orange, monomers 1-106), and TAD E (blue, monomers 107-307) are recovered by random loops (green arrows) according to the connectivity *ξ* and persistent long-range connectors (red bars), corresponding to peaks of the 5C data. **B.** Three realizations of the polymer model. **C** Encounter frequency matrix of the simulated polymer model, showing two TADs, where the off-diagonal points correspond to fixed connectors. **D.** Comparison between *β* computed from experiments and simulations data, confirming that the present polymer model accounts for the statistics of encounter frequencies.

We started with a Rouse chain (Fig. 4A (gray)) and added connectors between monomer pairs corresponding to peaks of the EP matrix (red). Three polymer realizations, simulated with the two types of connectors, are shown in Fig. 4B, characterized by a radius of Gyration *R*_*g*_ = 6.4. We computed the EF-matrix (Fig. 4C) that showed several similarity with the experimental data (compare Fig. 1A with Fig. 4B), for which two TAD-like structures are visible. Next, we quantified the similarity between the two matrices by comparing the decay exponent β_*n*_,(*n* = 1‥307). We used the function 12 to fit the EP of monomers 1 − 307 after long time polymer simulations (see Materials and Methods). The fitted value for *β* shows an excellent agreement with the experimental *β* values (Fig 3C). To conclude, long deterministic and short-range stochastic interactions are sufficient to account for the decay rate of EP extracted from coarse-grained 5C data.

### 3.5 Simulation of transient property of TADs: conditional encounter probabilities and mean times for three genomic sites to interact

We showed previously how to construct a polymer, the statistical properties of which match the ones extracted from 5C data. However, these data cannot be used to study transient properties of the chromatin, as they represent a static genomic encounter interactions, averaged over cell realizations. We shall use now the reconstructed polymer described above, constrained by the steady-state properties of the 5C data (Fig. 5A), to evaluate the transient properties of the chromatin. We estimate the mean first encounter time and the probability that monomer 26 (position of the Linx) meet monomer 87 (Xite) before monomer 64 (Chic1). These monomers represents three key sites on the X chromosome [5, 3], located in TAD D. We show three realizations and indicate the location of the three sites inside TAD D (yellow) and E (blue), that do not mix.

**Figure 5:**
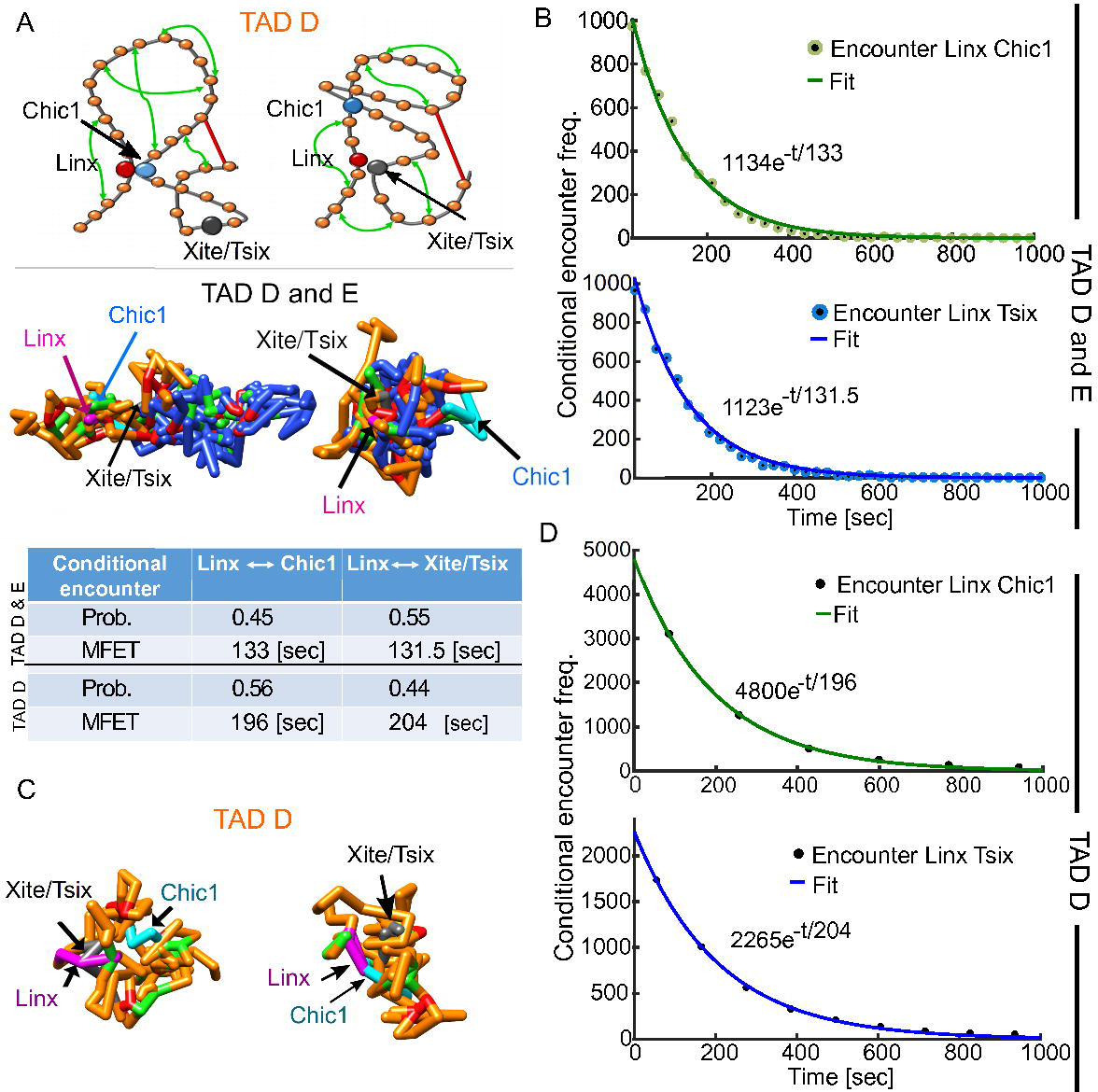
Transient properties of the chromatin: Conditional mean time and probability for three sites to meet. **A.** (upper panel) Representation of the polymer model for TAD D (orange, monomers 1-106), where loci Linx (monomer 26, red) meets Chic1 (monomer 67, cyan) and Xite/Tsix (monomer 87, gray), respectively. Random connectors (green arrows) and specific long range-connectors (red bar) are added, following the connectivity*ξ* recovered from data. Fixed connectors (red bars) correspond to specific peaks of the 5C data. Three realizations (bottom panel) of the polymer model containing TAD D and E, show the encounter of Linx (magenta) with Chic1 (cyan), and Xite/Tsix (gray), respectively. The color code is from the upper panel. **B.** Histogram of the conditional encounter times between Linx and Chic1 (upper panel, green),and Linx and Xite/Tsix (bottom panel, blue) with TAD D and E. **C.** Two polymer realization with a single TAD D (monomers 1-106, orange), showing the encounter between Linx (magenta) and Xite/Tsix (gray, left panel), and the encounter between Linx and Chic1 (cyan, right panel). **D.** Histogram of the conditional encounter times for a polymer with only TAD D, showing an exponential decay as in sub-figure C.

We started the polymer simulation from the steady-state distribution and use 10 000 runs. As predicted by the narrow escape theory [27, 28], the encounter time between two of the three monomers is Poissonian, as confirmed by their computed distribution (Fig. 5B), for which the reciprocal of the encounter rate is the mean encounter time. We found that the EP is *P* = 0.55, while the mean times are quite comparable of the order of 131s (see table C in Fig. 5). Finally, to check the impact of TAD E on the mean encounter time inside TAD D, we ran another set of stochastic simulations after we removed TAD E (Fig. 5D). Surprisingly, the EP was inverted compared to the case of no deletion, while the mean encounter has increased by almost 50% to 195s (Linx to Chic1) and 205s (Linx to Xite). This result suggests that TAD E contributed in modulating the interaction probability and the mean time, and thus further indicates that the search time inside a TAD depends on neighboring chromatin configuration.

## 4 Discussion

We presented here a general method to construct a coarse-grained polymer model from 5C encounter probability (EP) matrix. This construction preserves some of the statistical properties of the 5C data, such as the decay rate of the EP of each monomer. The method uses Rouse polymer as a starting point, where persistent long-range connectors, corresponding to local peaks of the 5C data, and random connectors are added in each realization. Connectors are represented by springs between long-range monomer-pairs and reflect the ensemble of chromatin architectures, as seen in the 5C-matrix. Sub-regions with encounter enrichment (TADs) are accounted for by adding connectors between random monomer-pairs, the number of connectors was extracted from EP-matrix. Finally, although contact maps (5C data) represents a steady-state distribution of an ensemble of looping events, patterns such as TADs appear in a large sample of millions of nuclei. We have demonstrated that using polymer reconstruction, transient properties can be studied, such as the mean encounter time between any two sites. The present approach is quite different from reconstruction methods that consist on inferring 3D structures of a genome from 5C data from contact frequency between sequences, which can be assumed to be Poissonnian [29] or not [30, 31].

Interestingly, TADs emerge here as a consequence of adding independent connectors randomly attached to monomers inside the subpart of the polymer that defines the TAD. These connectors could represent binding proteins such as cohesin [19] in the heterogeneous chromatin population of the 5C experimental data. Previous models explore the effect of connectors between regions of the chromatin [14, 13] and examine the consequence on the EP-decay rate. Here, we use random connectors to resolve a reverse engineering problem, and to recover the degree of connectivities from the EP-decay rate (see Fig.2). To accurately account for the statistical properties of the EP-matrix, we used the exponent of each monomer, and estimated the strength of interactions. For that reason, the present model extends the binders model developed in [14] that consider constant binding.

The mean looping time in free space depends only on the distance between monomers, while in confined domain this time also includes the effect of the boundary, which affects distant monomers [32, 28, 26]. It was thus surprising that a chromatin TAD subregion could influence the mean encounter between monomers located in a different region (Fig. 5). This effect is certainly due to the long-range inter-TAD interactions. Finally, the present method and algorithm can be used to reconstruct a polymer model at a given scale (number of monomers and number of bps coarse-grained in a monomer) that should be specified by the user. The polymer model following the specification described here, should reproduce the local condensation and can be used to study any transient chromatin properties involving any monomer pair.

## 6 Acknowledgments

We thank E. Heard, A. Amitai and L. Giorgetti for many interesting discussions on this subject.

